# Alternative catalytic residues in the active site of Esco acetyltransferases

**DOI:** 10.1101/766543

**Authors:** Tahereh Ajam, Inessa De, Nikolai Petkau, Gabriela Whelan, Vladimir Pena, Gregor Eichele

## Abstract

Cohesin is a protein complex encircles the DNA and regulates the separation of sister chromatids during cell division. Following a catalytic mechanism that is insufficiently understood, Esco1 and Esco2 acetyltransferases acetylate Smc3 subunit of cohesin, thereby inducing a stabilization of cohesin on DNA. As a prerequisite for structure-guided investigation of enzymatic activity, we determine here the crystal structure of the mouse Esco2/CoA complex at 1.8 Å resolution. We reconstitute the entire cohesin as a tetrameric assembly and use it as a physiologically-relevant substrate for enzymatic assays *in vitro*. Furthermore, we employ cell-based complementation studies in mouse embryonic fibroblast deficient for *Esco1* and *Esco2*, as a means to identify catalytically-important residues *in vivo*. These analyses demonstrate that D567/S566 and E491/S527, located on opposite sides of the MmEsco2 active site cleft, are critical for catalysis. Our experiments supports a catalytic mechanism of acetylation where residues D567 and E491 are general bases that deprotonate the ε-amino group of lysine substrate, via two nearby serine residues - S566 and S527-that possess a proton relay function.

## Introduction

Cohesin is a ring-like protein complex that is composed of the four subunits Smc1, Smc3, Scc1/Rad21, and Scc3/SA. Smc1 and Smc3 belong to the family of SMC (structural maintenance of chromosomes) proteins, which contain ATPase head domains. Cohesin topologically traps chromatin inside its ring structure, functioning as a molecular connector that ensures equal segregation of sister chromatids during mitosis ^1-4^. Additionally, cohesin facilitates postreplicative homologous recombination repair of double-strand DNA breaks, and regulates gene expression *via* the formation of chromatin loops ^5-9^. Cohesin is loaded onto chromatin by the Scc2-Scc4 loader complex and released by Wapl and Pds5 ^10-15^. Both cohesin loading and unloading depend on the ATPase activity of the Smc head domain ^10,16-19^.

Once loaded, DNA interacts with a basic patch residing on the Smc3 head domain and thereby stimulates its ATPase activity. A notable structural feature of this basic patch is the presence of two neighboring conserved lysines ^19,20^. Acetylation of these residues by yeast acetyltransferase Eco1 or its mammalian orthologues Esco1 and Esco2 (<u>es</u>tablishment of <u>co</u>hesion) decreases the positive charge of the patch, which weakens DNA binding and lessens ATPase activity ^19^. This in turn counteracts the activity of the release factors Wapl-Pds5. As a result, Esco activity stabilizes cohesin on DNA ^21^. In vertebrates, cohesion establishment additionally involves Sororin, which competes with Wapl for binding to Pds5 and in this way counteracts the releasing activity of Wapl-Pds5 ^22,23^.

Esco1 and Esco2 belong to the GNAT (GCN5-related N-acetyltransferase) family. These two isozymes consist of divergent N-termini, followed by a C2H2 zinc finger and a conserved C-terminal acetyltransferase domain ^24^. Esco1 and Esco2 differ in several respects. Esco1 is evenly expressed throughout the cell cycle, while Esco2 is highly abundant during the S-phase ^25,26^. Esco1 but not Esco2 directly interacts with cohesin *via* Pds5 ^27^. Esco2 interacts with the replication proteins, PCNA (proliferating cell nuclear antigen) ^28,29^ and MCM (minichromosome maintenance protein complex) ^30,31^. Esco1 mutation is associated with endometrial cancer ^32^ and mutations in Esco2 cause RBS (Roberts syndrome), a congenital disease ^33-35^. In RBS metaphase chromosomes show a loss of cohesion in the PCH (pericentric heterochromatin) while cohesion is maintained in the arms ^36^. A significant fraction of *Esco1*-deficient mice is viable (this study), while *Esco2*-deficient mice always die early in development ^26^.

An acetyltransferase typically employs a general base to deprotonate the ε-amino group of a substrate lysine which initiates acetyl transfer from AcCoA. Intriguingly, crystal structure of human acetyltransferase HsESCO1, reported by Kouznetsova *et al*., led the authors to propose that the general base is provided by the substrate rather than the enzyme. Thus, D107 of SMC3 might deprotonate K105 and K106 of SMC3 ^37^. In contrast, Rivera-Colon *et al*. proposed that D810 of HsESCO1 plays the role of a general base. Indeed, mutating this aspartate to asparagine abrogates acetylation of an Smc3 peptide mimicking SMC substrate ^38^. A subsequent study by Chao *et al*. has observed that in the *Xenopus* xEco2/Smc3 peptide structure, the Smc3 D107 does not point towards the ϵ-amino group of the substrate lysines but interacts with two conserved R621 and W623 residues of xEco2. This suggests that D107 of Smc3 plays a role tethering the enzyme to the substrate, rather than acting as a general base ^39^. In agreement with Rivera-Colon *et al*., Chao *et al*. propose that D677 (the equivalent of D810 of HsESCO1) could serve as a general base. However, both studies noted that this particular aspartic acid is not strictly conserved among Esco homologs, suggesting that other residues at the active site may also contribute to catalysis.

To shed light into the catalytic mechanism of Esco1 and Esco2, we determine the crystal structure of murine Esco2 in complex with CoA at atomic resolution, establish a robust framework for assessing enzymatic activity *in vitro* and *in vivo*, and perform mutagenic analysis of the active center. An alternative catalytic model is proposed.

## Results

### Structure of MmEsco2^368-592^ in complex with Coenzyme A

We crystallized MmEsco2^368-592^ protein consisting of the C2H2 zinc finger and the acetyltransferase domain. Crystals diffracted to 1.8 Å resolution. By making use of the natively bound zinc ion, the crystal structure was determined by SAD (single-wavelength anomalous dispersion) from a dataset collected at the zinc peak wavelength (Supplementary Table S1). The refined MmEsco2^368-592^ structure revealed continuous electron density, except for two short, structurally disordered and hence unresolved regions (residues 368-383 and 501-514, Fig. 1A). The zinc finger (residues 385-416), is located N-terminally of the catalytic domain, and consists of two β-strands and one α-helix that encircle the zinc ion (Fig. 1B). Residues 423-592 of MmEsco2^368-592^ reveal an overall fold similar to that of other GNAT family acetyltransferases ^40^. This fold consists of a conserved catalytic domain (β5, β6, β7, α3, and α4), which is flanked by structurally variable regions (Fig. 1B). The CoA cofactor is natively present in a complex with MmEsco2 in a groove formed by β7 and β8 strands and α3 and α4 helices (Fig. 1B). This is reminiscent of the position of the CoA or AcCoA in other GNAT family members ^40^.

**Figure 1.**
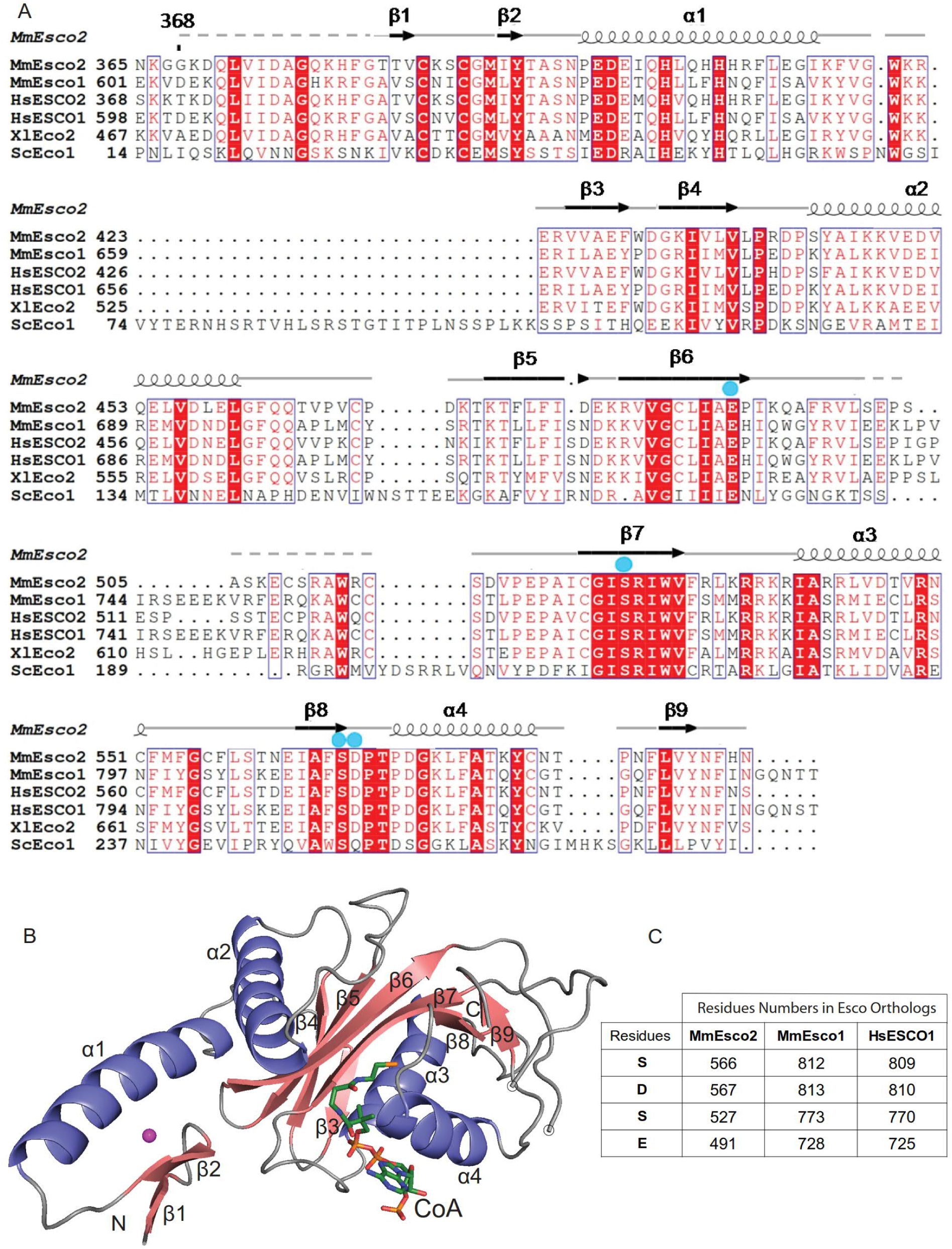
Structure of the MmEsco2^368-592^/CoA complex. (**A**) Sequence alignment ^51^ of Esco orthologs. Sequences shown are *Mus musculus* MmEsco1, MmEsco2, *Homo sapiens* HsESCO1, HsESCO2, *Xenopus laevis* XlEco2 and *S. cerevisiae* ScEco1. Invariant residues are shown with a red background, and highly conserved residues are boxed. Numbering and secondary structural elements above the sequence alignment are for the MmEsco2^368-592^ sequence. Dashed lines mark the disordered regions. Blue circles indicate residues that might be involved in the abstraction of the proton from the ε-amino group of the substrate lysine. (**B**) Ribbon representation of the MmEsco2^368-592^/CoA complex. α-helices are shown in blue, β-strands in raspberry, and loop regions in grey. CoA is represented as sticks and colored according to elements: carbon, green; nitrogen, blue; sulfur, orange; oxygen, red and the zinc ion shown as a magenta sphere. There is an unresolved region in a loop connecting β6 and β7. Start and end point of this region is indicated by empty circles. (**C**) Numbering of equivalent putative catalytic residues of MmEsco2 in MmEsco1 and HsESCO1 sequences.

### The active site architecture of MmEsco2^368-592^ and identification of candidate catalytic residues

We searched for residues in the active site cleft of MmEsco2^368-592^, which could act as a general base for catalyzing the nucleophilic attack of the lysine ε-amino group on the AcCoA thioester bond. Structural superposition with xEco2 in complex with a Smc3 peptide conjugated with CoA at K105 ^39^, enabled identification of candidate catalytic residues in MmEsco2, with reference to the ε-amino group of K105. The most obvious candidate residue is D567 that may act in conjunction with S566; the latter potentially acting as a proton relay. Noteworthy, the equivalent D810 was previously suggested as general base in HsESCO1 ^38^ (for an overview of residues equivalence, see Fig. 1C). S566 and D567 are located at the CoA binding pocket, S566 in the C-terminus of the β8 strand and D567 in a flexible loop connecting β8 strand and α4 helix (Fig. 2A). The γ-oxygen of S566 and δ-oxygen of D567 are ∼ 5 and ∼ 4.4 Å away from the ε-amino group of K105 (Fig. 2A). The distance of the δ-oxygen of D567 and the γ-oxygen of S566 is 5.7 Å, consistent with a proton relay function. We also considered S527 as a possible relay, with its γ-oxygen being ∼ 8.7 Å away from ε-amino group of K105 (Fig. 2A). E491 could serve as general base. Its ε-oxygen of is ∼ 9.8 Å away from ε-amino group of K105. Notably, a water molecule located in between these four residues and CoA, similar to the one observed in the xEco2/K106-CoA structure ^39^, might be involved in proton transfer (Fig. 2B). E491, S527 and S566 residues are invariant among Esco homologs. D567 residue is also highly conserved except for yeast Eco1 where this residue is a glutamine (Fig. 1A).

**Figure 2.**
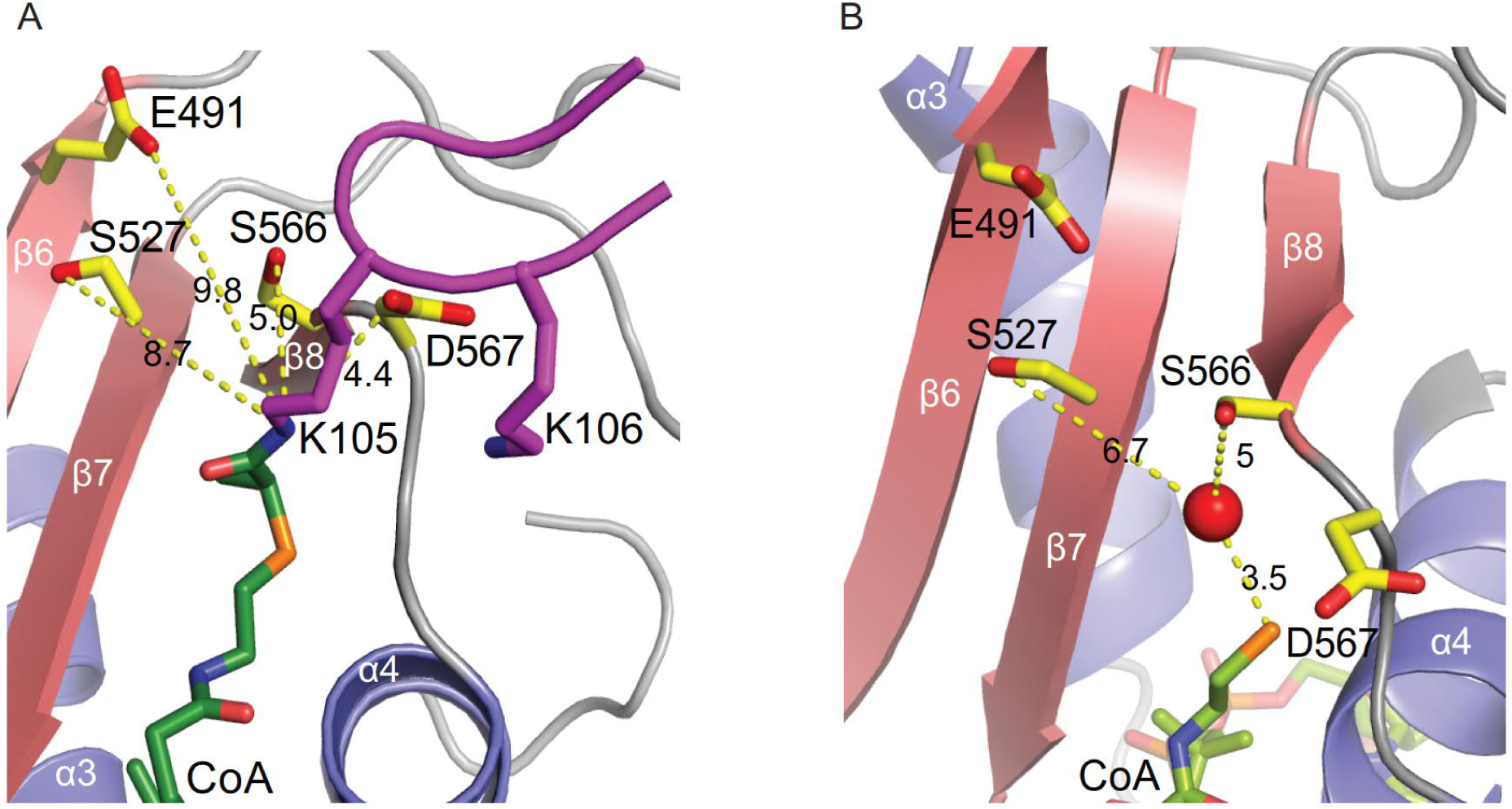
The active site of the MmEsco2^368-592^/CoA complex. (**A**) Close-up view of the active site of the MmEsco2^368-592^/K105-CoA model. The position of the K105 was modeled by superposition with the structure of xEco2 in complex with a K105-CoA peptide [PDB ID code 5N1W]. Side chains of putative catalytic residues are shown. Dashed lines indicate the distances in angstrom from relevant atoms of the putative catalytic residues to the ε-amino group of K105. Smc3 peptide is shown in magenta. (**B**) A water molecule is located in the active site of MmEsco2^368-592^/CoA between the candidate catalytic residues and CoA.

### Establishing of an acetylation assay that mimic the physiological conditions

To investigate whether the selected candidate residues contribute to catalysis, we mutated S566 and/or D567 and measured catalytic activities with trimeric or tetrameric cohesin ring substrates. Expression of full-length MmEsco1 and MmEsco2 wild type proteins gave very low yields, but recombinant full-length HsESCO1 was readily expressed in Sf9 cells (Supplementary Fig. S1A). Of note, the structure of the active site of MmEsco2 and HsESCO1 are very similar (Supplementary Fig. S2A). Thus, we applied HsESCO1 for the following reconstitution-based assays.

A major drawback of the enzymatic assays reported before for Esco proteins is that either autoacetylation or acetylation of peptide mimetics were used, rather than physiological substrates. Recombinant cohesin assembled as a trimer was also employed, although it does not thoroughly represent physiological substrate ^41^. To enhance the relevance and accuracy of our observations, we therefore decided to reconstitute entire cohesin rings recombinant and use them for enzymatic assays.

We first generated human trimeric (Supplementary Fig. S1B) and tetrameric recombinant cohesin as previously described ^41^. Trimeric and tetrameric substrates were acetylated by HsESCO1, at K105 and K106 of SMC3 in the presence of ATP, circular, linear or relaxed DNA (Figs. 3A and 3B). To detect SMC3 acetylation, we used a monoclonal antibody that specifically recognizes Smc3 singly acetylated on K106 or doubly acetylated on K105 and K106 ^23^. Acetylation was not detected in reconstituted cohesin substrate when ATP hydrolysis was inhibited ^41^ (Supplementary Fig. S1C) or DNA was absent (Figs. 3A and 3B). Importantly, tetrameric substrate was much more efficiently acetylated than trimeric cohesin (Figs. 3C and 3D). We conclude that in presence of ATP and DNA, HsESCO1 efficiently acetylates cohesin rings. Given that cohesin rings are efficiently acetylated only in the tetrameric form, and that acetylation is dependent of ATP and DNA, we conclude that the assay that we established recapitulates to a good extent the expected occurrence of HsEsco1 acetylation under physiological conditions.

**Figure 3.**
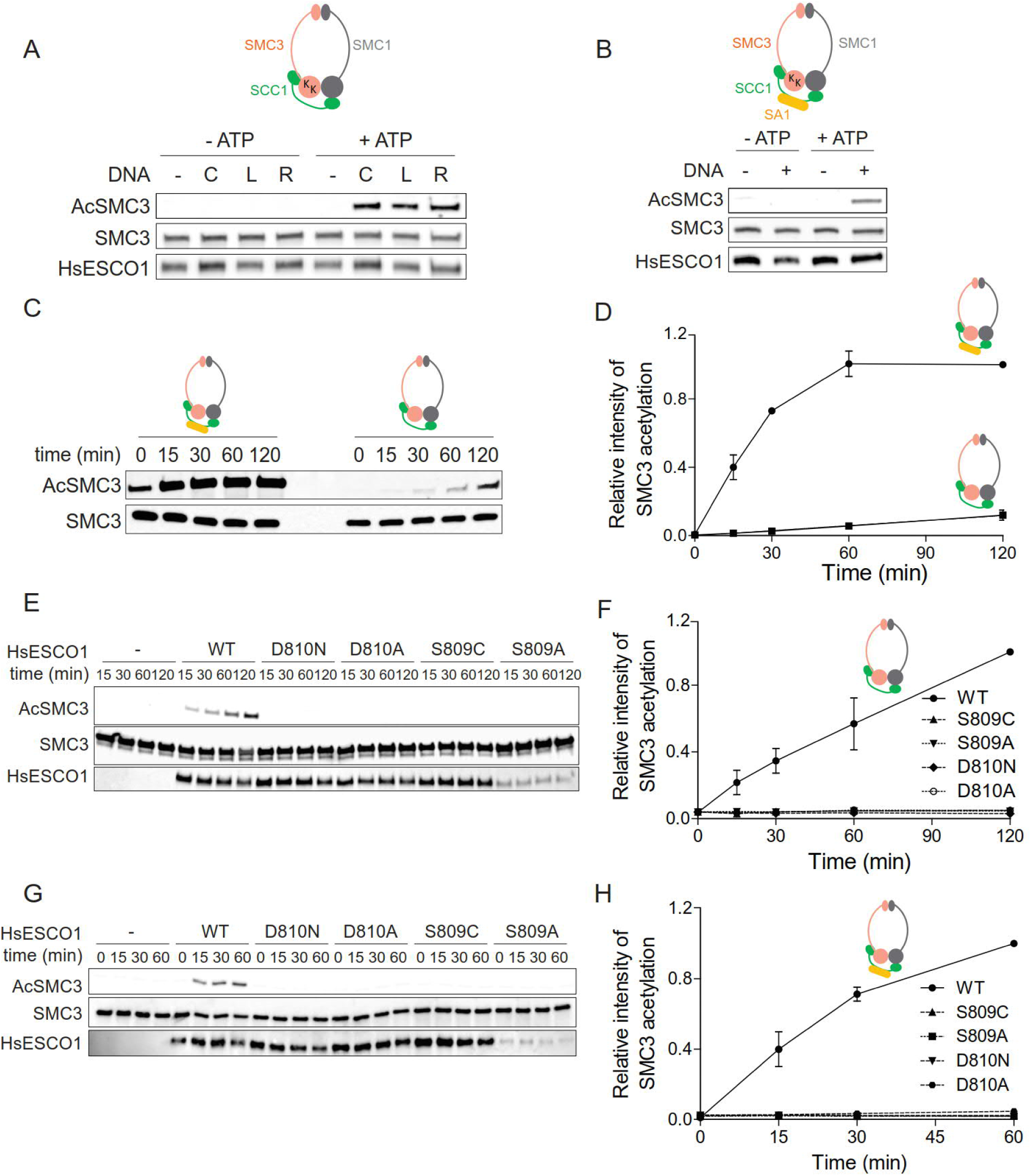
S809 and D810 are required for HsESCO1 activity to acetylate recombinant cohesin substrates. (**A**) The purified recombinant trimeric cohesin complex was incubated with HsESCO1 and AcCoA in the presence or absence of ATP and different topologies of the pcDNA3 plasmid (linear DNA [L], relaxed circular DNA [R] or covalently closed circular DNA [C]). (**B**) The purified tetrameric cohesin complex was incubated with HsESCO1 and AcCoA in the presence or absence of ATP and covalently closed circular DNA. (**C**) and (**D**) Time course of SMC3 acetylation after incubation of purified trimeric or tetrameric cohesin complexes with HsESCO1 in the presence of ATP, DNA and AcCoA. Half-times for the tetramer is ∼ 20 min. Acetylation of the trimer is considerably slower. (**E**) and (**F**) Time course of SMC3 acetylation after incubation of purified trimeric cohesin with wild-type or mutants of HsESCO1. (**G**) and (**H**) Time course of SMC3 acetylation after incubation of purified tetrameric cohesin with wild-type or mutants of HsESCO1. All data were normalized to maximum signal and are shown as mean ± SEM (n=2).

### Mutations in putative catalytic residues affect acetyltransferase activity of Esco1 in reconstituted *in vitro* assay

Having the mimicking physiological reconstituted system at hand, we next quantified the rate of SMC3 acetylation for site-directed mutants of residues S809 and D810. Recombinant mutant proteins HsESCO1^S809A^, HsESCO1^S809C^, HsESCO1^D810A^, and HsESCO1^D810N^ were produced at similar levels as wild-type controls, except for HsESCO1^S809A^ which gave a lower yield. Recombinant proteins were incubated with cohesin tri- and tetrameric complexes. Wild-type HsESCO1 efficiently acetylated both substrates (Figs. 3E, 3F, 3G and 3H). Remarkably, none of the mutant proteins acetylate cohesin trimers or tetramers (Figs. 3E, 3F, 3G, and 3H). Rivera-Colon *et al*. ^38^reported that wild-type protein and the four mutants used here show similar thermal stability, arguing against the possibility that mutant HsESCO1 proteins were unfolded and hence became catalytically inactive. Together, these results indicate that D810 and the neighboring S809 are required for catalytic activity of the HsESCO1 enzyme, when using reconstituted recombinant cohesin substrates. We infer from the HsESCO1 experiments that S566 and D567 of MmEsco2 are catalytic residues.

### Mutations in putative catalytic residues affect acetyltransferase activity of Esco1 and Esco2 in cells

The existing interactions between several accessory proteins and the tetrameric cohesin ring rises the possibility that the former affect the geometry of the active site pocket and the way substrates bind. Therefore, we asked whether the serine and aspartic acid mutants, completely inactive in the above reconstitution assays, are also catalytically inactive when assayed in MEFs. These cell-based complementation assays involve transfection of full-length wild-type or mutant *MmEsco1/2* into MEFs lacking either MmEsco1 or MmEsco2.

To obtain *Esco1*-deficient MEFs (MEFs^*Esco1-/-*^), an *Esco1* conditional knockout mouse was generated using the Cre/LoxP system allowing deletion of exons 2 and 3 (Supplementary Figs. S3A-S3D). MEFs^*Esco1-/-*^ were arrested in G1 (Fig. 4A), showed a strong reduction in Smc3 acetylation compared to controls (Fig. 4B). Residual Smc3 acetylation is likely due to MmEsco2 that is present at low-amounts during the G1-phase ^25,26^. Transfection of MEFs^*Esco1-/-*^ with *MmEsco1-myc* restored strong Smc3 acetylation (Fig. 4B). Next, we transiently transfected MEFs^*Esco1-/-*^ with single residue mutants that were catalytically inactive in the *in vitro* reconstitution experiments (Figs. 3E-3H). It should be noted that all MmEsco1 mutants tested were chromatin bound (Supplementary Fig. S4C). Transfection of MEFs^*Esco1-/-*^ with *MmEsco1*^*S812A*^, *MmEsco1*^*S812C*^, and *MmEsco1*^*D813A*^ showed significant acetyltransferase activity, reaching up to 30 to 50% of wild-type level in contrast to the reconstitution-based assays (Figs. 4C and 4D). Note that mutant and wild-type cells were equally arrested in G1 (Supplementary Fig. S4A).

**Figure 4.**
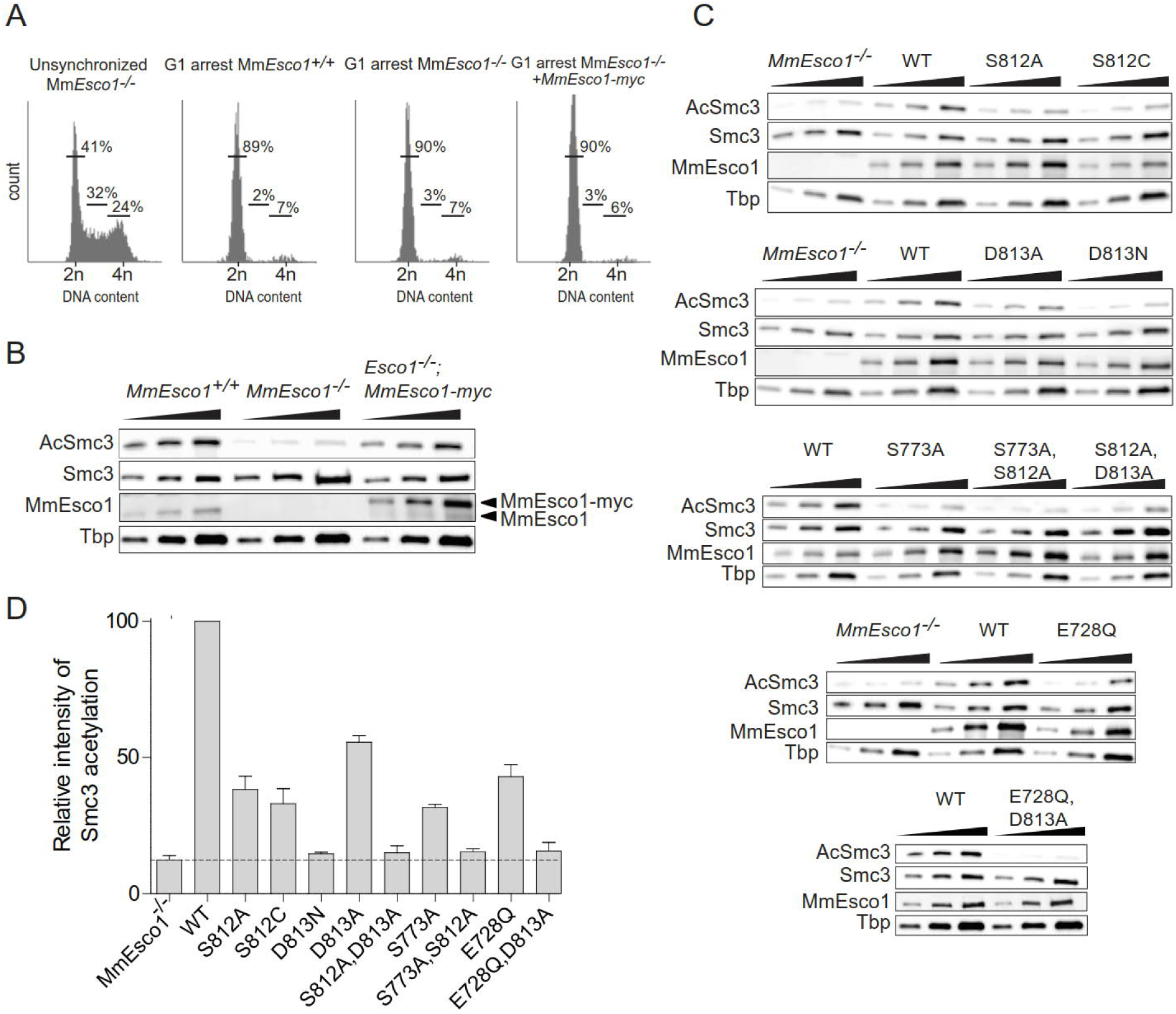
S812, D813, S773, and E728 function cooperatively in the catalysis of MmEsco1. (**A**) Flow cytometry profiles of G1-phase arrested MEFs^*Esco1+/+*^, MEFs^*Esco1-/-*^ and MEFs^*Esco1-/-*^ expressing ectopic wild-type MmEsco1. Unsynchronized MEFs^*Esco1-/-*^ were used as reference. The numbers show the percentage of cells in G1, S, G2/M phase. (**B**) Immunoblot of MEFs^*Esco1+/+*^, MEFs^*Esco1-/-*^ and MEFs^*Esco1-/-*^ expressing ectopic wild-type MmEsco1, arrested in G1-phase. Of note, transiently transfected MmEsco1 was expressed at a level of about 4-fold that of endogenous MmEsco1. (**C**) Immunoblots of MEFs^*Esco1-/-*^ transiently expressing wild-type or mutant versions of MmEsco1, arrested in G1-phase (Supplementary Fig. S4A). Note that MEFs^*Esco1−/−*^ expressed comparable levels of different MmEsco1 variants. (**D**) Quantification of the data shown in C. The dotted line indicates the Smc3 acetylation value for MEFs^*Esco1-/-*^. Data were normalized to wild-type signal and are shown as mean ± SEM (n=3). In all immunoblotting, the chromatin-bound fractions were analyzed using a 2-fold serial dilution.

Rivera-Colon *et al*. showed that HsESCO1^D813N^ does not acetylate SMC3 peptide substrate ^38^, which is consistent with our reconstitution experiments with cohesin ring substrates (Figs. 3F and 3H). When we transfected MEFs^*Esco1-/-*^ with MmEsco1^D813N^ Smc3 acetylation was completely abolished. This strong effect of D813N mutation is likely caused by the loss of the general base character, as well as of the interference of asparagine with the positioning of the ε-amino group of the substrate lysines.

Of note, D813A still exhibits considerable activity (Figs. 4C and 4D), suggesting that E728 (Figs. 1C and 2A) takes over the role of a general base. Consistent with a mechanism that involves a second general base residue is the observation that the double mutant protein E728Q;D813A resulted in base-line level activity (Figs. 4C and 4D). As the active site geometry places the carboxyl group of E728 almost 10 Å away from the ε-amino group of K105, S812 and S773 might act as proton shuttles. Consistently, we find that S812A;S773A double mutants are nearly inactive (Figs. 4C and 4D).

So far, we identified several MmEsco1 residues that mediate Smc3 acetylation in cells. We next examined whether the corresponding residues are also involved in catalysis in MmEsco2. Wild-type, *Esco2*^*-/-* 26^, and *Esco2*^*-/-;Esco2-myc*^ MEFs were arrested in S-phase (Fig. 5A) at which time endogenous MmEsco2 is known to be maximally expressed ^25,26^. Consistent with previous published data^26^, MEFs^*Esco2-/-*^ showed a 60% reduction in Smc3 acetylation compared to controls (Figs. 5B and 5D). Residual Smc3 acetylation is most likely due to MmEsco1, which is expressed throughout the cell cycle^25,26^. Transfection of MEFs^*Esco2-/-*^ with *MmEsco2-myc* restored Smc3 acetylation to wild-type levels (Figs. 5B and 5D). Next we stably transfected MEFs^*Esco2-/-*^ with corresponding Esco2 mutants, which were all chromatin bound (Supplementary Fig. S4D). Reminiscent of the experiments done with MmEsco1 mutants, single mutants *MmEsco2*^*S566A*^, *MmEsco2*^*D567A*^, *MmEsco2*^*S527A*^ and *MmEsco2*^*E491Q*^ were still able to acetylate Smc3 to some extent (Figs. 5C and 5D). This was not the case with *MmEsco2*^*D567N*^ and the double mutants *MmEsco2*^*S566A;D567A*^ and *MmEsco2*^*S527A;S566A*^ (Figs. 5C and 5D; for synchronization of mutant cells see Supplementary Fig. S4B).

**Figure 5.**
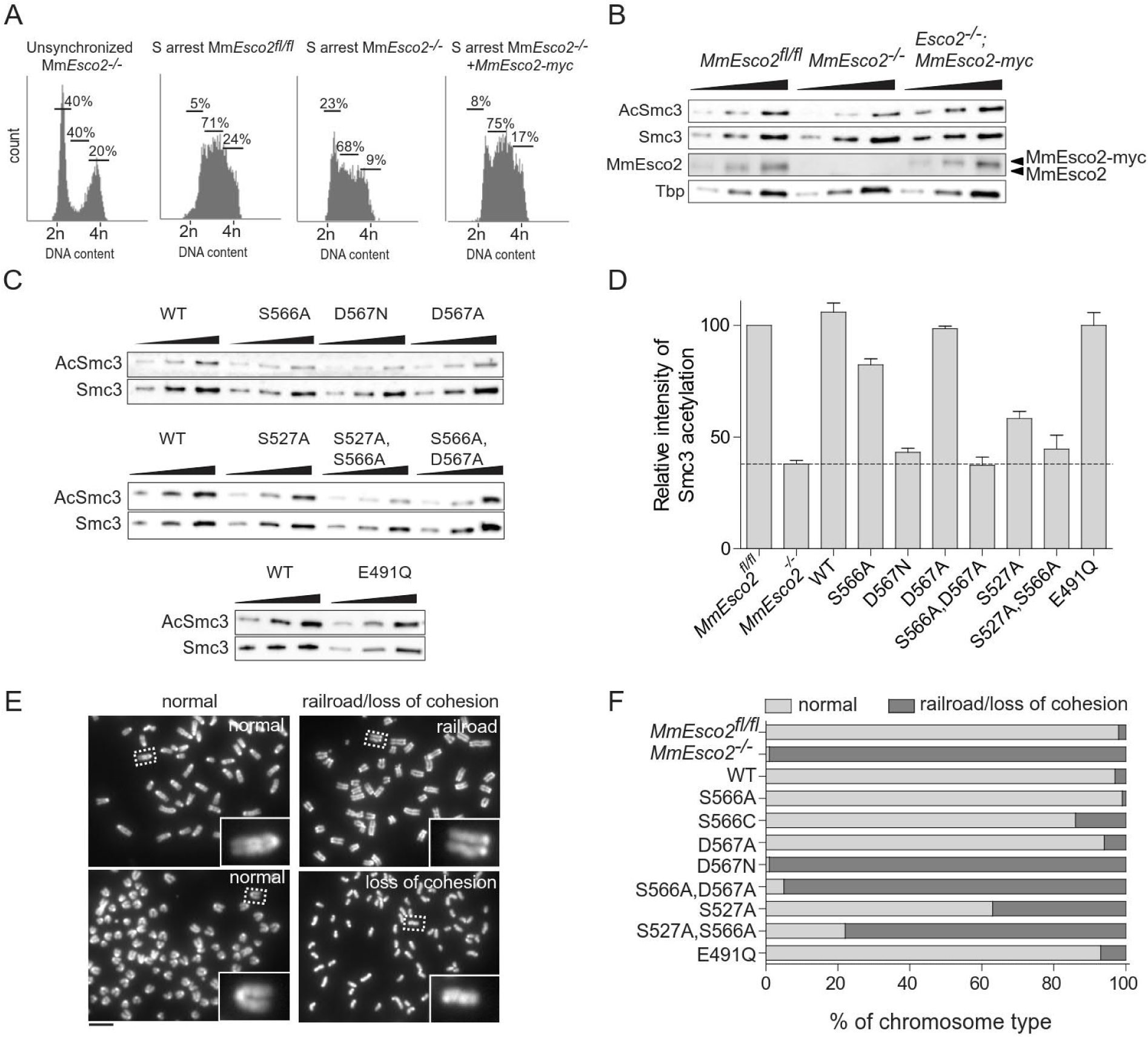
S566, D567 and S527 function cooperatively in the catalysis of MmEsco2. (**A**) Flow cytometry profiles of S-phase arrested MEFs^*Esco2fl/fl*^, MEFs^*Esco2-/-*^ and MEFs^*Esco2-/-*^ expressing ectopic wild-type MmEsco2. Unsynchronized MEFs^*Esco2-/-*^ were used as reference. The numbers show the percentage of cells in G1, S, G2/M phase. (**B**) Immunoblot of MEFs^*Esco2fl/fl*^, MEFs^*Esco2-/-*^ and MEFs^*Esco2-/-*^ expressing ectopic wild-type MmEsco2, arrested in S-phase. Of note, stably transfected MmEsco2 was expressed at a similar level as the endogenous MmEsco2. (**C**) Immunoblot of MEFs^*Esco2-/-*^ stably expressing wild-type or mutant versions of MmEsco2, arrested in S-phase (Supplementary Fig. S4B). Note that MEFs^*Esco2-/-*^ express comparable levels of Esco2 variants (see Supplementary Fig. S4D). (**D**) Quantification of the data shown in C. The dotted line indicates the Smc3 acetylation level seen in MEFs^*Esco2-/-*^. Data were normalized to signal of MEFs^*Esco2fl/fl*^ and are shown as mean ± SEM (n = 3). In all immunoblotting experiments, the chromatin-bound fractions were analyzed using a 2-fold serial dilution. (**E**) Representative prometaphase chromosomes spreads including normal, railroad and single chromatids. Scale bar: 10 µm. (**F**) Frequency of chromosome types were assessed in 1000 prometaphase chromosomes.

While these results are in close agreement with the effect of the same mutants on MmEsco1 activity, the effect of single mutations (S566A, D567A, S527A and E491Q) on Smc3 acetylation was slightly less pronounced than that observed for MmEsco1. For instance, MmEsco2^S566A^ and MmEsco2^S527A^ showed up to 60-80% residual acetyltransferase activity relative to wild-type (Figs. 5C and 5D). In the case of Esco1, the residual activity was in the 30-50% range. In addition, MmEsco2^D567A^ and MmEsco2E492Q single mutants acetylated Smc3 to the same extent as its wild-type control (Figs. 5C and 5D). These differences between Esco1 and Esco2 might be caused by the different chromatin context in which the two enzymes act.

In summary, our data indicate that both MmEsco1 and MmEsco2 engage alternative catalytic residues, supporting a shared mechanism of acetylation, in spite of the distinct biological contexts where the two enzymes function.

### Effect of catalytic mutants of MmEsco2 on sister chromatid cohesion

RBS patients are characterized by loss of function mutations in HsESCO2 causing sister chromatid cohesion defects ^34,35^. Such defects are also seen in *Esco2*-deficient MEFs (Fig. 5E; ^26^). We examined which of the MmEsco2 mutants lead to a cohesion defect phenotype. MEFs^*Esco2-/-*^ were stably transfected with wild-type or different mutant versions of *MmEsco2*, and were synchronized in prometaphase, isolated by mitotic shake-off, and used for chromosome spread preparations. Consistent with the Smc3 acetylation experiments (Figs. 5C and 5D), chromosome morphology analysis revealed that the single mutants MmEsco2^S566A^, MmEsco2^D567A^, MmEsco2^S527A^, and MmEsco2^E491Q^ that still enable acetyltransferase activity (Figs. 5C and 5D) restore wild-type appearance of cohesion, to a good extent (Fig. 5F). However, catalytically inactive single or double mutants MmEsco2^D567N^, MmEsco2^S566A;D567A^ and MmEsco2^S527A;S566A^, exhibited strong sister chromatid cohesion defects (Fig. 5F). As would be expected, the sister chromatid cohesion assay points to the same catalytic residues as the acetylation assays.

## Discussion

The initial step of a lysine acetyltransferase reaction is deprotonation of the ε-amino group of substrate lysine, mediated by a general base, most commonly an aspartate or a glutamate. Subsequently, the deprotonated ε-amino group carries out a nucleophilic attack on the carbonyl carbon of enzyme-bound AcCoA. This eventually results in the transfer of the acetyl moiety from AcCoA to the substrate lysine.

The active site cleft of Esco acetyltransferases has two walls. One consists of α-helices 3 and 4 whose side chains position AcCoA and a loop that connects a short β-strand (β8) to α-helix 4. The opposite wall consists of β-strands 5, 6 and 7 (Figs. 1A and 1B). Candidate catalytic residues protrude from the walls (Fig. 2A). Previous attempts to identify catalytic residues by mutational analyses of Esco1 and Esco2 homologs did not arrive at the same results. In the case of HsESCO1 an aspartate residue belonging to the substrate has been proposed to act as a general base ^37^. However, in a more recent study ^38^, this role was attributed to an aspartate (D810) from the active site of HsESCO1. Mutation of this highly conserved aspartate to an asparagine inactivated the enzyme, according to an enzymatic assay that made use of Smc3 peptide as a substrate. Crystal structure of xEco2, the Xenopus homolog of Esco1 and Esco2, complexed with an isolated peptide of the Smc3 substrate supports the catalytic role of this aspartate as a general base ^39^.

Our own structure-function studies of MmEsco2 also identified this particular aspartate (D567 in MmEsco2, Fig. 6), located on the loop between β8 and α4, as a general base. A D567N mutation was enzymatically inactive in reactions using recombinant trimeric and tetrameric cohesins as substrates. Furthermore, this particular mutant failed to acetylate Smc3 in cells and establish sister chromatid cohesion.

**Figure 6.**
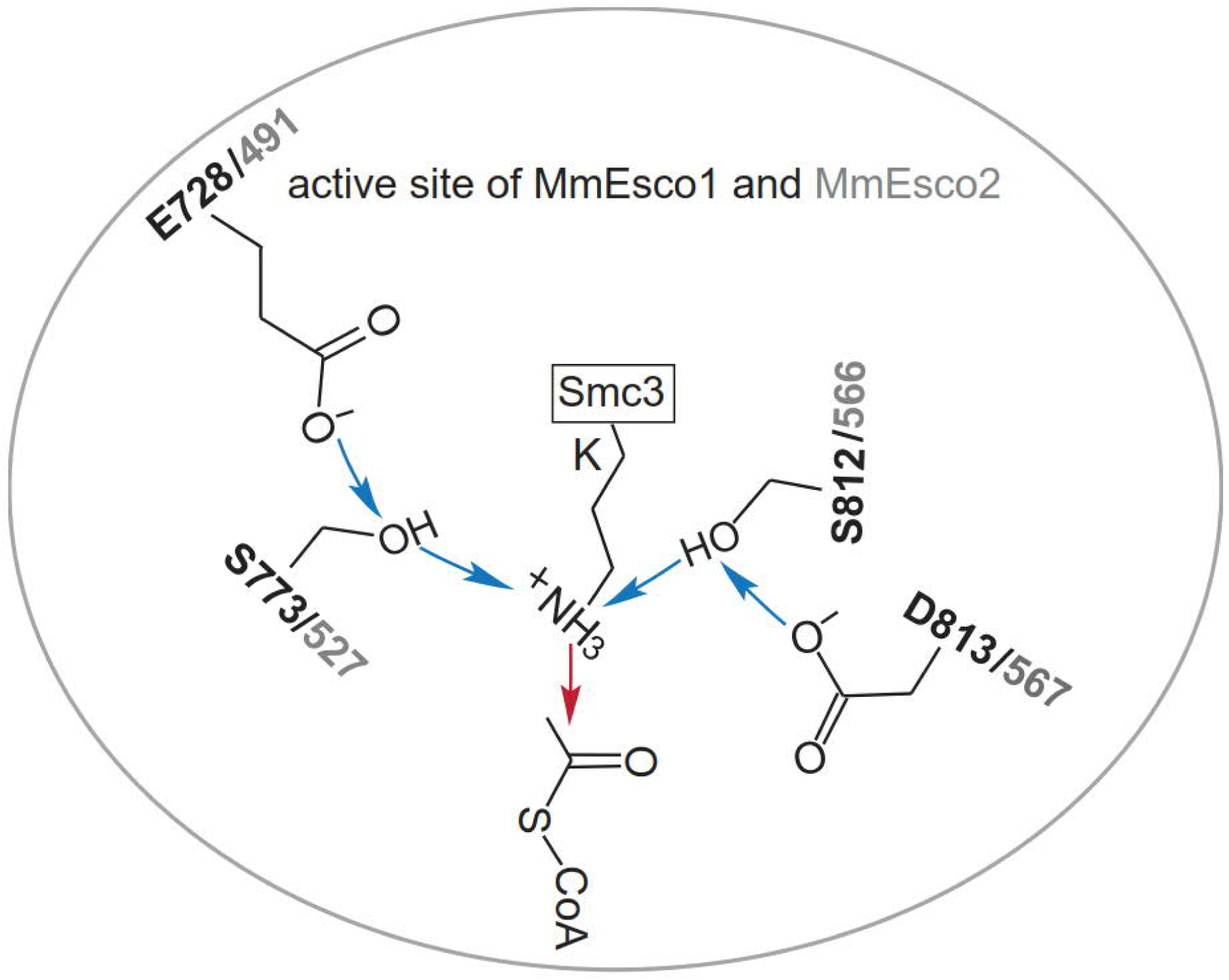
Proposed catalytic mechanism for Esco1 and Esco2. The proposed mechanism involves D566 and/or E491, acting as a general base, to initiate the reaction by abstracting a proton from the hydroxyl group of S566 and/or S527. Subsequently, the hydroxylate of S566 and/or S527 can then act as base catalyst to deprotonate the amino group of K105 and/or K106 (located on a hairpin in Smc3 structure ^52^). Subsequently, the nucleophilic attack of the amine on the carbonyl carbon of AcCoA occurs. Blue and red arrows indicate deprotonation and the nucleophilic attack, respectively. The numbering of putative catalytic residues is based on the MmEsco2 sequence (see Fig. 1C for residue equivalence).

Surprisingly, although mutation of D567 to an alanine abolished acetylation *in vitro*, the enzyme showed appreciable activity *in vivo*, as observed in two different cell-based assays. This indicates that the role of D567 as a general base is taken over by another residue. A possible candidate is E491, located on the β-strand 6 at the opposite of the CoA binding pocket (Fig. 2A). Mutation of the equivalent residue from HsESCO1 to a glutamine was previously shown to abolish acetylation of an SMC3 peptide ^38^. Notably, this mutation reduces activity by 50% in cell-based assays, while the double mutant E491Q; D567A is completely inactive in cell-based assays. These observations support the view that both of these residues from the active site cleft act as general bases.

Serine residues, S527 and S566, are located in close proximity to E491 and D567, respectively (Fig. 2A). Mutation of S566 to either a cysteine or an alanine abolishes acetylation *in vitro* (Figs. 3E-3H). However, both mutants were partially active in cell-based assays (Figs. 4C and 4D). Crystal structure of MmEsco2 carrying the S566C mutation is virtually identical with the wild-type protein (data not shown), suggesting that substitution of this residue impairs catalysis rather than the structure. Mutation of the second serine residue, such as S527A is known to abolish acetylation *in vitro* ^38^. This is consistent with our observation that S527A reduces substantially acetyltransferase activity in cell-based assays (Figs. 4C and 4D). Taken together, these experiments argue for roles of S527 and S566 in catalysis, possibly acting as a proton relay for E491 and D567 (Fig. 6). A proton relay function of serine has been proposed for the Dat acetyltransferase in which a glutamate residue deprotonates the substrate amino group through a serine residue ^42^. Whether a nearby water molecule (Fig. 2B) is also involved in proton transfer is as yet unclear. In support of this idea is the finding that a water molecule is found at the same position in xEco2 ^39^.

Our work highlights the crucial importance of designing and employing suitable enzymatic assays, when complex substrates such as cohesin are involved. Firstly, we observed *in vitro* that the presence of SA1 in the composition of tetrameric cohesin (therefore a closer mimic of the physiological context) enables a notably more efficient acetylation than the trimeric version (Figs. 3C and 3D). Furthermore, mutations that abolish enzymatic activity versus this tetrameric cohesin substrate (e.g. S566A, S566C and D567A) still enable acetylation of cohesin *in vivo*, where even more factors are associated to cohesin. Although the residual activity is relatively weak, in one case can rich up to 50% of the wild-type activity (Figs. 4C and 4D). This is indicative for redundancy mechanisms that are at work *in vivo*, owing to additional cohesin regulators that are absent *in vitro*, such as Pds5A, Pds5B, MCM and chromatin ^22,27,31^.

Furthermore, we noticed the identical mutations in either MmEsco1 or MmEsco2 evoke Smc3 acetylation deficiencies to a similar qualitative extent (Figs 4D and 5D). Nonetheless, there are some quantitative differences. This difference between Esco1 and Esco2 might reflect the fact that they act *in vivo* in a very different molecular context and do not have identical binding partners.

Taken together, we propose that two general bases present in Esco enzymes abstract the substrate proton *via* serine residues (Fig. 6). Presence of the catalytic residues D567 and S566 in the CoA binding wall of the active site cleft and E491 and S527 protruding from the wall made of β-strands 5, 6 and 7 suggest that the active sites of Esco1 and Esco2 could be mirror symmetrical. Esco1 and Esco2 sequentially acetylate two distinct, neighboring lysines located in the basic patch of Smc3 ^39^. An interesting possibility is that the “duplication” of the catalytic residues has evolved as a mechanism required for acetylation of both substrate lysine residues, without the need of dissociation and re-association of the enzyme-substrate complex.

## Material and Methods

### MmEsco2^368-592^ cloning, expression and purification

Truncated MmEsco2^368-592^ with a C-terminal His-tag was cloned into the pFL vector ^43^. The expression construct was transformed into DH10MultiBacY cells and recombinant bacmid was purified with a QIAprep Spin Miniprep Kit (Qiagen). V_0_ and V_1_ virus generation was performed using SF9 cells as previously described ^44^. Hi5 cells were infected with V_1_ virus for protein expression. Cells were harvested after 48 h and resuspended in lysis buffer (50 mM HEPES pH 7.2, 200 mM NaCl, 10% glycerol, 2 mM DTT, and complete EDTA-free protease inhibitors cocktail [Roche]) and lysed using a microfluidizer. The lysate was cleared by centrifugation and the supernatant was applied onto a 50 ml anion-exchange Q-Sepharose column (GE Healthcare) equilibrated with lysis buffer. The bound proteins were eluted with a linear gradient from 100 mM to 1 M NaCl. The peak fractions containing Esco2 were pooled and applied onto a 1 ml Ni-NTA Superflow column (Qiagen), equilibrated with 20 mM HEPES pH 7.2, 500 mM NaCl, 10% glycerol, 2 mM DTT, and 10 mM imidazole. The bound proteins were eluted with a linear imidazole gradient of 15-250 mM. The peak fractions were pooled, concentrated to a volume of 2 ml and applied onto a S75 16/600 pg size exclusion column (GE Healthcare), equilibrated with 10 mM HEPES pH 7.2, 150 mM KCl, 5% glycerol and 2 mM DTT. Peak fractions were concentrated and stored at −80 °C.

### MmEsco2^368-592^ crystallization and structure determination

MmEsco2^368-592^ was crystallized using the sitting-drop vapor-diffusion method at 20°C. Crystals were obtained from droplets consisting of 100 nl of MmEsco2^368-592^ (10 mg/ml) and 100 nl crystallization solution (100 mM Tris, 20% (v/v) 2-Methyl-2, 4-pentanediol [MPD], pH 8). After harvesting, crystals were cryoprotected in 15–20% ethylene glycol and flash frozen in liquid nitrogen.

Diffraction data were collected at beamline PXII of SLS (Paul Scherrer Institute, Villigen, Switzerland), processed and scaled using XDS ^45^. By making use of the natively bound zinc ion, the crystal structure was determined by single-wavelength anomalous dispersion (SAD) from a 2.3 Å dataset collected at the zinc peak wavelength (Native I in the Supplementary Table S1). The initial model was then refined against a dataset of higher resolution 1.77 Å (Native II in the Supplementary Table S1). The final model was built manually using COOT32 ^46^ and structure refinement was performed with Phenix33 ^47^.

### Human ESCO1 cloning, expression and purification

HsESCO1 was cloned into the pFastbac-HTC vector (Invitrogen) with an N-terminal His-tag. HsESCO1 variants was expressed in Sf9 cells. Cells were harvested after 48-72 h and resuspended in lysis buffer (20 mM HEPES pH 7.5, 300 mM NaCl, 10% glycerol, 30 mM imidazole, 1 mM TCEP and complete EDTA-free protease inhibitors cocktail [Roche]) and lysed by sonication. The lysate was cleared by centrifugation. Subsequently, the supernatant was incubated with equilibrated Ni-NTA beads (Qiagen) for 2 h at 4 °C. Bound proteins were washed with high salt buffer (lysis buffer containing 1 M NaCl). Proteins bound to Ni-NTA beads were eluted with lysis buffer containing 150 mM NaCl, 500 mM Imidazole and dialyzed for 16 h against dialysis buffer (20 mM HEPES pH 7.5, 100 mM NaCl, 10% glycerol, 1 mM DTT). The dialyzed samples were snap-frozen.

### Human trimeric and tetrameric cohesin complex cloning, expression and purification

To produce trimer complex, HsScc1 was cloned into a 438-C vector (Addgene, 55220), containing an N-terminal His-tag followed by a maltose binding protein (MBP) tag and a tobacco etch virus (TEV) protease cleavage site. Cloning into this vector was achieved by the ligation independent cloning (LIC) method. HsSmc3-FLAG and HsSmc1-His in pFastbac and combined Smc1, Smc3-FLAG, Scc1 and His-SA1 in a pFL multibac vector were provided by Jan-Michael Peters lab (Research Institute of Molecular Pathology, Vienna).

The trimer cohesin complex was expressed in Sf9 cells using coinfection with Smc1-His, Smc3-FLAG, and Scc1-MBP viruses. Cells were lysed in lysis buffer (20 mM HEPES pH 7.5, 500 mM NaCl, 10% glycerol, 1 mM DTT and complete EDTA-free protease inhibitors cocktail [Roche]) supplemented with 0.02% NP40 and 1 mM PMSF. After sonication and clarification by centrifugation, the lysate was applied onto a 5 ml amylose column (GE Healthcare) equilibrated with lysis buffer. The bound proteins were eluted with a linear gradient of 10-100 mM maltose. The peak fractions were pooled, concentrated and applied on a S200 16/600 pg size exclusion column (GE Healthcare), equilibrated with 10 mM HEPES pH 7.2, 150 mM KCl, 5% glycerol and 2 mM DTT. Peak fractions were concentrated and snap-frozen.

The tetramer cohesin complex was expressed in Hi5 cells using coinfection with Smc1, Smc3-FLAG, Scc1, and HIS-SA1 viruses. Cells were lysed in lysis buffer (50 mM HEPES pH 7.5, 300 mM NaCl, 10% glycerol, 2 mM DTT, 30 mM imidazole and complete EDTA-free protease inhibitors cocktail [Roche]) supplemented with 1 mM TCEP, 1 mM Pefabloc (Sigma) and 0.05% Tween-20. After sonication and clarification by centrifugation, the supernatant containing tetrameric cohesin was filtered using a 0.8 μm filter (Millipore). Subsequently, the lysate was incubated with 1 ml of Ni-NTA beads for 2 h at 4 °C. Ni-NTA beads were washed with 10 beads volume (BV) of lysis buffer, followed by 10 BV of high salt buffer (lysis buffer containing 1 M NaCl), lysis buffer and finally low salt buffer (lysis buffer containing 150 mM NaCl). Proteins bound to Ni-NTA beads were eluted with lysis buffer containing 150 mM NaCl and 250 mM imidazole. Eluate was incubated with 200 µl anti-FLAGM2 agarose beads (Sigma) for 2 h at 4 °C. The complex was eluted with elution buffer (25 mM HEPES pH 7.5, 150 mM NaCl, 10% glycerol, 1 mM DTT and 0.5 mg ml-1 FLAG peptide) and snap-frozen.

### Site-directed Mutagenesis

Point mutations in HsESCO1, MmEsco1 and MmEsco2 were introduced with the QuikChange II XL site-directed mutagenesis kit (Agilent Technologies) according to the manufacturer’s manual and were verified by DNA sequencing.

### *In vitro* acetylation assay

Acetylation assays were performed by preincubation of 500 nM of trimer (to compare acetylation of tetrameric and trimeric cohesin by HsESCO1, 100 nM we used) or 100 nM of tetramer with 240 µM ATP, 10 µM AcCoA, 3.3 nM pcDNA3.1 plasmid, 25 mM HEPES pH 7.5, 25 mM NaCl, 1 mM MgCl_2_, and 0.05 mg ml^-1^ BSA at 32 °C. After 1 h, 50 nM HsESCO1 and additional NaCl (100 mM final concentration) were added and incubated at 37 °C. The reactions were stopped at the different time points by adding an equal volume of 2X SDS loading buffer, and denatured at 100 °C for 5 min.

### Generation of *Esco1* conditional knockout mouse line

The *Esco1* targeting vector, containing a long (7.3 kb in size) and a short (3.8 kb in size) homology arm, two loxP sites flanking exons 2 and 3 and a FRT-flanked neomycin cassette, was generated by Polygene. After Validation of the targeting construct, we linearized the vector by NotI restriction endonuclease and electroporated it into the 129/SvPas embryonic stem (ES) cells. Subsequently, neomycin-resistant clones were screened by Southern blotting. The verified ES cell clones were injected into C57BL/6J blastocysts. Resulting chimeric animals were mated with ACTB-FLPe mice ubiquitously expressing FLP-recombinase ^48^. Removal of the neomycin cassette and germ line transmission were validated by PCR. To generate *Esco1*^*-/-*^ mice for subsequent MEF isolation, *Esco1*^*fl/fl*^ mice were crossed to a ubiquitous UBC-Cre driver mouse line^49^. For Southern blotting, the probe (361 bp) was amplified with primers TCTCGTCATTTCAGAAACCATC and GCTCACCTATGCTCACATGAAG. For PCR genotyping, a combination of three primers was used (P1: CACCACACTGGCATTAAGTTCTAGG, P2: CCTTACAGTGATGAAACTGAGCACC, P3: GAGTTTCCTGTAGCCAGAGGATGG).

### Cell culture, transfection, synchronization, extract preparation and immunoblotting

Wild-type and indicated mutants of MmEsco1 were cloned into pEF6/Myc-His B vector (Thermo Fisher Scientific) containing C-terminal myc and His tags. Immortalized MEFs^*Esco1-/-*^ cultured in standard medium (DMEM, supplemented with 10% fetal bovine serum [FBS], 100 U/ml penicillin and 100 µg/ml streptomycin) were transiently transfected with the wild-type and mutant versions of *MmEsco1*. To synchronize cells in G1, 36 h after transfection, the medium was changed to DMEM medium supplemented with 10% FBS and 25 µM lovastatin (Thermo Fisher Scientific). Cells were harvested after 24 h and synchronization was assessed by flow cytometry.

Wild-type and mutants *MmEsco2-myc/his* and *H2B-mCherry* were cloned into the pVITRO2-hygro-mcs vector (InvivoGen) in two steps. First, full-length *MmEsco2* was cloned into the pcDNA3.1/myc-His vector (Thermo Fisher Scientific). Subsequently, *MmEsco2-myc/his* and *H2B-mCherry* were amplified from the vectors pcDNA3.1/myc-His and pcDNA3-*H2B-mCherry* (Addgene, 20972), respectively, and cloned into the pVITRO2-hygro-mcs vector. Immortalized MEFs^*Esco2fl/fl*^ were grown to confluence in standard medium and transduced with Ad-Cre-GFP adenoviruses (SignaGen) in low serum medium (DMEM, supplemented with 3% FBS). After two days, the medium was changed to fresh low-serum medium and cells were cultured for another 48 h. Immortalized MEFs^*Esco2-/-*^ were stably transfected with wild-type and mutant versions of Esco2. Clones were then screened using a plate reader by measuring mCherry fluorescence levels. Subsequently, clones that stably expressed the mutant proteins close to the endogenous MmEsco2 level were selected using Western blotting. For synchronization, cells were treated twice with 2 mM thymidine for 14 h with an intermittent release of 9 h. Cells were harvested 2 h after second thymidine release and further processed for subsequent analyses.

For whole-cell extracts, cells were collected, washed in 1x cold PBS, resuspended in 2x SDS loading buffer and sonicated. Chromatin fractionation was performed according to the protocol described ^50^.

### Prometaphase chromosome spreads and immunofluorescence analysis

MEFs were grown in standard culture medium. Cells at 60% confluency were arrested using nocodazole (400 ng ml^-1^) for 4h. Mitotic cells were harvested by shaking off and incubated in 1 ml of 75 mM KCl for 20 min at 37 °C. Prometaphase chromosomes were fixed in methanol:acetic acid (3:1), dropped onto humidified positively charged microscope slides and visualized using DAPI staining.

### Antibodies

Rabbit antibody against MmEsco1 was generated using a haemocyanin-conjugated peptide comprising amino acids 521 to 606 of mouse Esco1 (1:1000). The following previously described custom-made antibodies were used: anti-Esco2 ^26^ (1:1000), mouse anti-acetyl-Smc3 (a gift from K. Shirahige) ^23^ (1:1000). The following commercial antibodies were used: rabbit anti-Smc3 (Cell Signaling D47B5, 1:3000), HRP-conjugated mouse anti-TBP (Abcam 197874, 1:5000), HRP-conjugated mouse anti-His tag (Novus 31055H, 1:1000).

## Author contributions

T.A., N.P., V.P. and G.E. designed the experiments. I.D. and V.P. collected the X-ray diffraction data and determined the crystal structure. T.A. conducted the experiments. T.A. and N.P. analyzed and interpreted the data. G.W. and T.A. generated the knock out mice. T.A. and G.E. wrote the paper. All authors reviewed and edited the manuscript.

## Acknowledgements

We thank S Mahsur, S Thiel, and J Wawrzinek for technical assistance. We are grateful to K Shirahige for providing Ac-Smc3 antibody; to JM Peters for sharing cohesin expression constructs and to P Cramer for sharing recombinant expression constructs. The research leading to these results has received funding from the Max Planck Society.

## Data availability

All data generated or analyzed during this study are included in the manuscript and supporting files.

## Supplementary Figure Legends

**Supplementary Figure S1.**
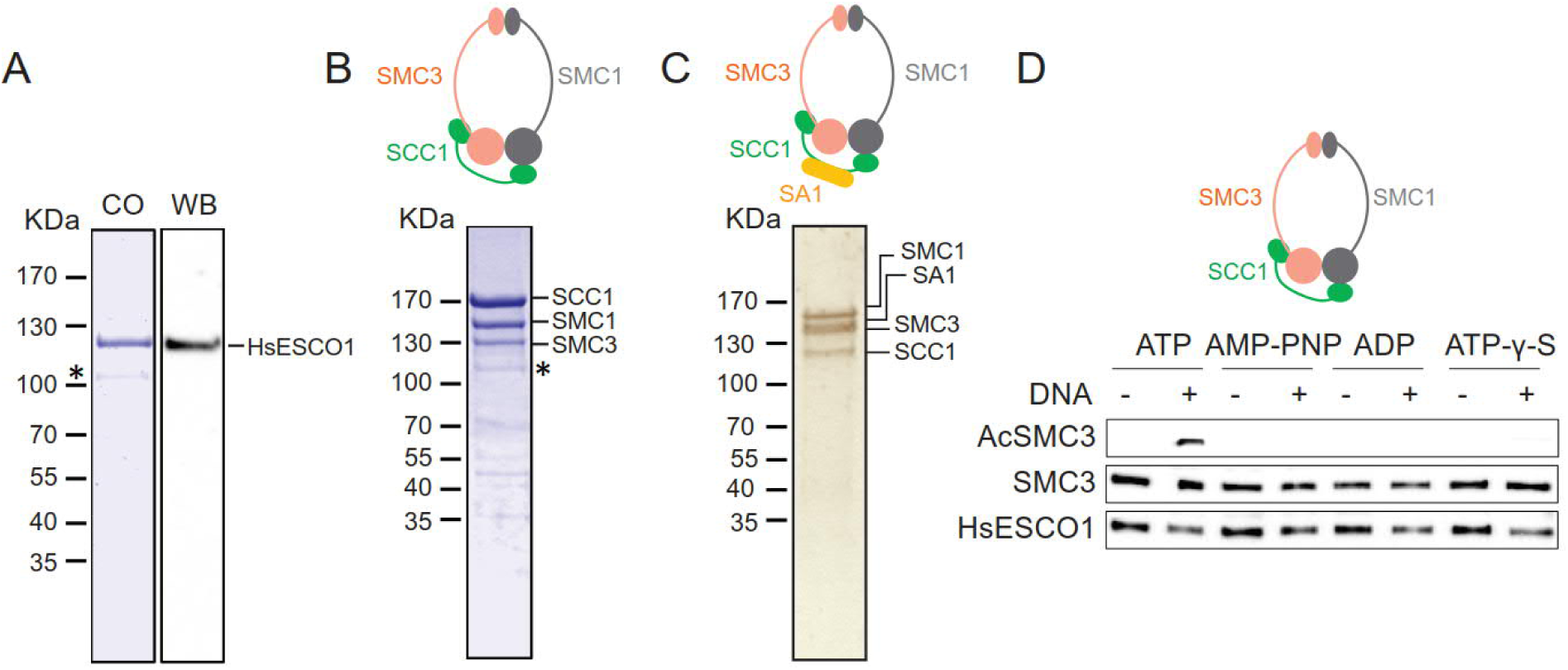
Purification and functional characterization of human Esco1 and cohesin complex. (**A**) Purified HsESCO1 analyzed by SDS-PAGE followed by Coomassie blue staining (CO) or Western blotting (WB). The protein was detected as a single protein band with a molecular mass of about 120 kDa. The asterisk indicates a minor contaminant. (**B**) Purified human trimeric cohesin analyzed by SDS-PAGE followed by Coomassie staining. The bands corresponding to SCC1, SMC1 and SMC3 are indicated on the right side. The masses of the molecular weight marker (kDa) are indicated on the left side. The asterisk indicates a minor contaminant. (**C**) The trimeric cohesin complex was incubated with HsESCO1 and AcCoA in the presence of ATP, adenylyl-imidodiphosphate (AMP-PNP), adenosine diphosphate (ADP) or adenosine 5′-[γ-thio] triphosphate (ATP-γ-S). The level of SMC3 acetylation level was analyzed by Western blotting.

**Supplementary Figure S2.**
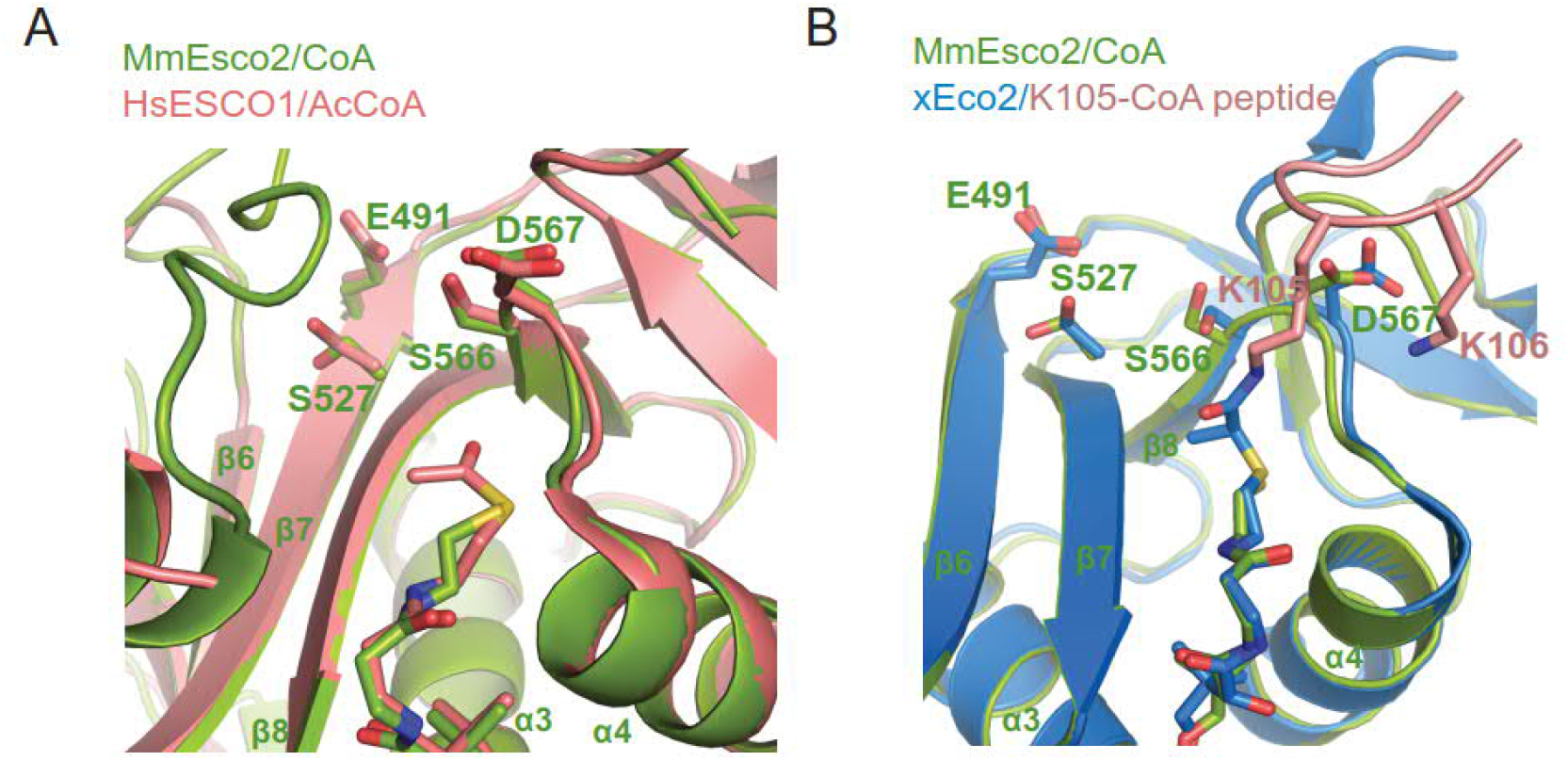
Active site of MmEsco2^368-592^/CoA. (**A**) Close-up view of the active sites of MmEsco2/CoA and HsESCO1/AcCoA. Putative catalytic residues of MmEsco2 (green) and HsESCO1 (raspberry; PDB ID code 4MXE) show high structural conservation. (**B**) Close-up view and comparison of the active sites of MmEsco2/CoA and xEco2/K105-CoA. Putative catalytic residues of MmEsco2 (green) and xEco2/K105-CoA (blue; PDB ID code 5N1W) are shown as sticks. K105 peptide Smc3 is shown in raspberry.

**Supplementary Figure S3.**
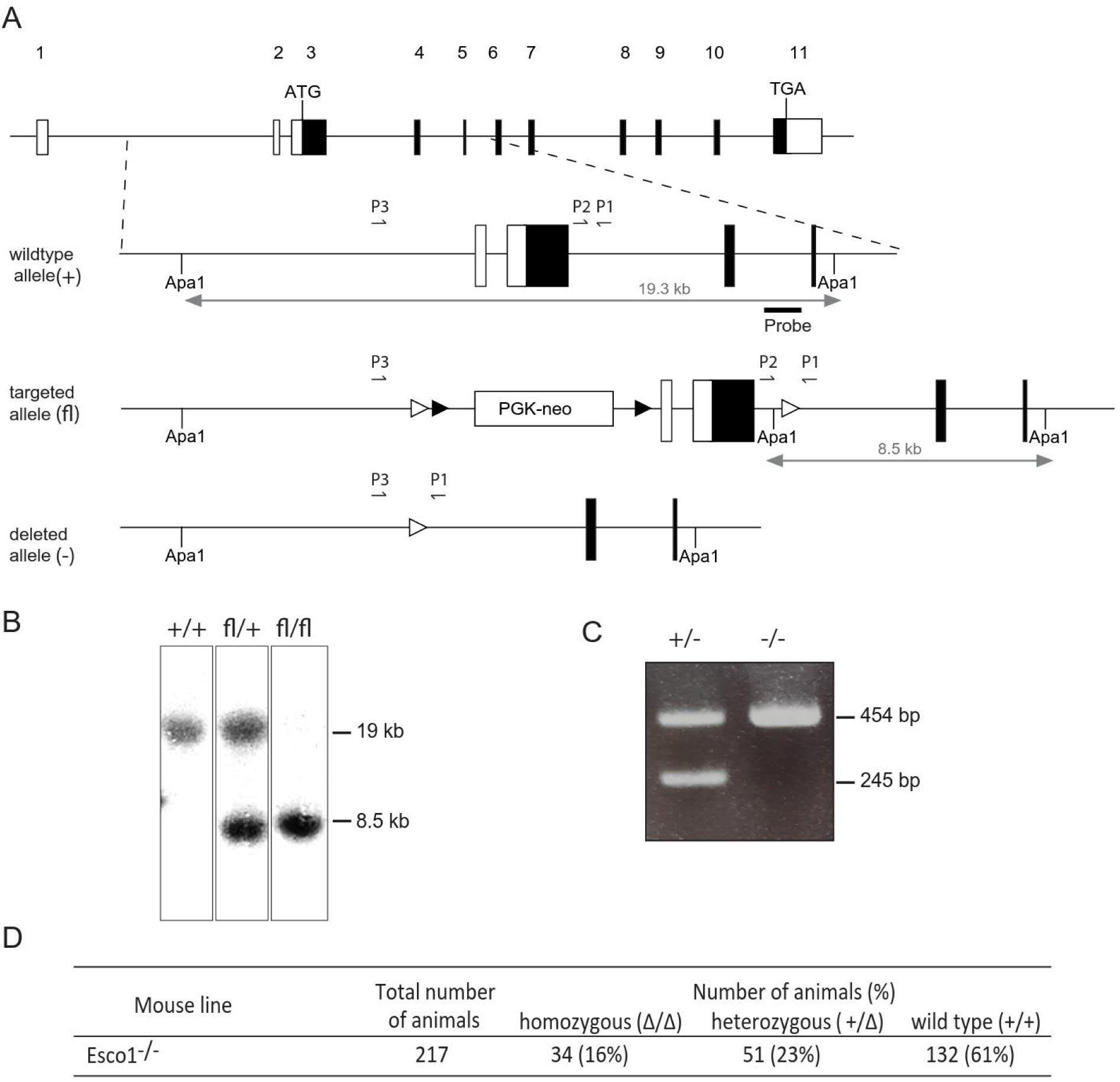
Generation of Esco1 conditional knockout mouse line. (**A**) Schematic representation of the *Esco1* wild-type locus (+), targeted allele (fl), and the null *Esco1* allele (-) created by Cre-mediated recombination. *LoxP* and *FRT* sites are marked by an empty or black triangle, respectively. (**B**) Southern blot of genomic DNA from mouse tails carrying the alleles indicated, digested with ApaI. The probe used is located between exons 4 and 5, as shown in A. (**C**) PCR genotyping of Esco1 alleles using the three primers indicated in A. The wild-type allele produces a 245-bp amplicon (primers P1, P2). Deletion of the LoxP-flanked region of the Esco1 locus leads to a 454-bp fragment (primers P1, P3). (**D**) Viability statistics of *Esco1*^*-/-*^ mouse line from heterozygous mating. Mice homozygously deficient for Esco1 are designated as *Esco1*^*-/-*^. Note that in *Esco1*^*-/-*^, the homozygous animals were born in a sub-Mendelian ratio, instead of 25 % only 16% of the newborn animals were homozygous.

**Supplementary Figure S4.**
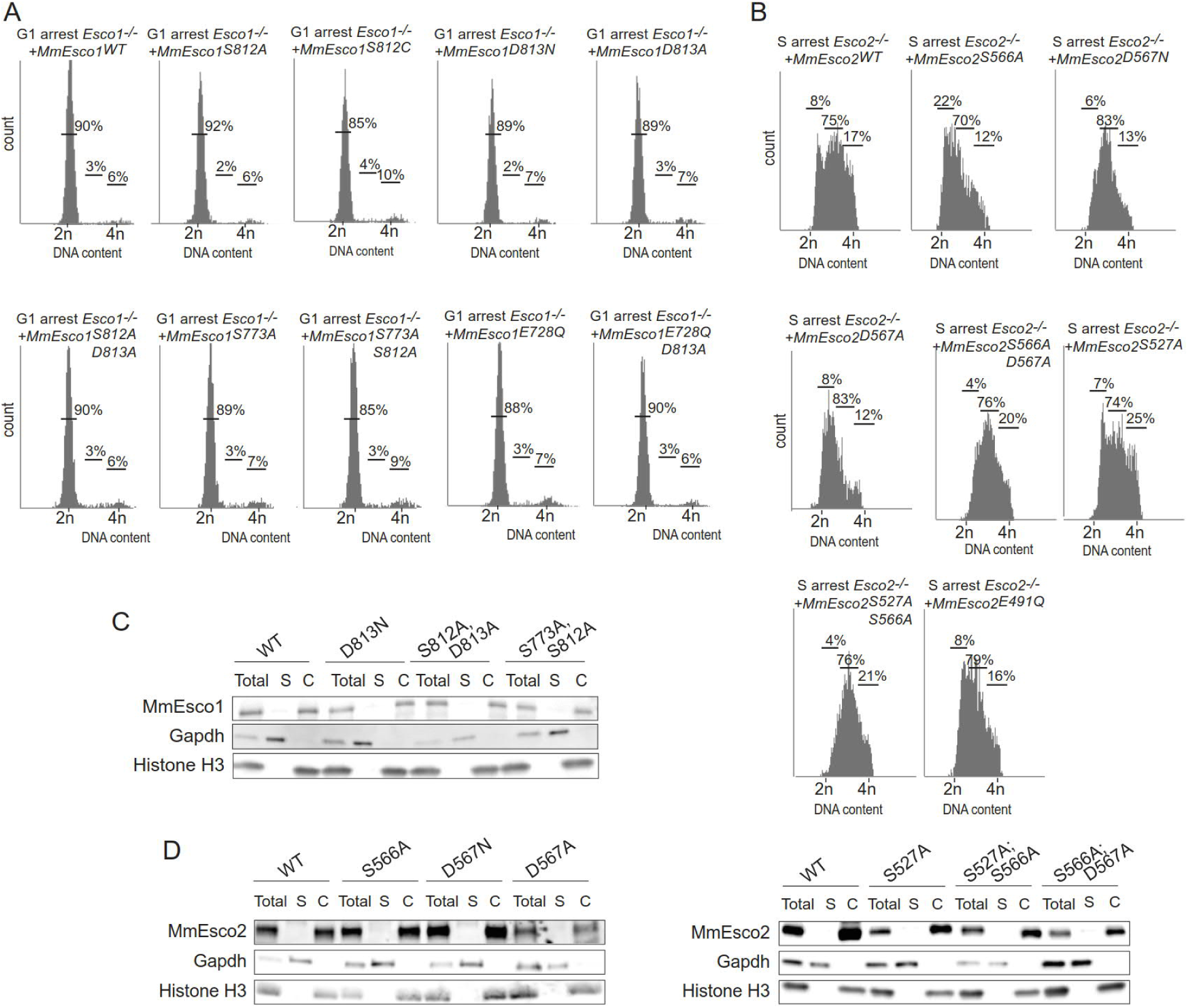
*In vivo* characterization of Esco1 and Esco2 mutants. (**A**) Flow cytometry profiles of G1-phase arrested MEFs^*Esco1-/-*^ expressing wild-type or mutant MmEsco1 (**B**) Flow cytometry profiles of S-phase arrested MEFs^*Esco2-/-*^ expressing wild-type or mutant MmEsco2. In A and B, the numbers show the percentage of cells in G1, S, G2/M phase. (**C**) Cell fractionation analysis of MEFs^*Esco1-/-*^ expressing wild-type and catalytically dead mutants of MmEsco1-myc. (**D**) Cell fractionation analysis of MEFs^*Esco2-/-*^ expressing equivalent amounts of ectopic MmEsco2-myc variants. In C and D, Total: total cell lysate S: soluble fraction, C: chromatin.

## Reference

1 Losada, A., Hirano, M. & Hirano, T. Identification of Xenopus SMC protein complexes required for sister chromatid cohesion. Gene Dev 12, 1986–1997, doi:DOI 10.1101/gad.12.13.1986 (1998).

2 Tanaka, T., Fuchs, J., Loidl, J. & Nasmyth, K. Cohesin ensures bipolar attachment of microtubules to sister centromeres and resists their precocious separation. Nat Cell Biol 2, 492–499, doi:10.1038/35019529 (2000).

3 Lammens, A., Schele, A. & Hopfner, K. P. Structural biochemistry of ATP-driven dimerization and DNA-stimulated activation of SMC ATPases. Current Biology 14, 1778–1782, doi:10.1016/j.cub.2004.09.044 (2004).

4 Uhlmann, F. SMC complexes: from DNA to chromosomes. Nat Rev Mol Cell Bio 17, 399–412, doi:10.1038/nrm.2016.30 (2016).

5 Peters, J. M., Tedeschi, A. & Schmitz, J. The cohesin complex and its roles in chromosome biology. Genes Dev 22, 3089–3114, doi:10.1101/gad.1724308 (2008).

6 Nasmyth, K. & Haering, C. H. Cohesin: Its Roles and Mechanisms. Annu Rev Genet 43, 525–558, doi:10.1146/annurev-genet-102108-134233 (2009).

7 Wendt, K. S. & Peters, J. M. How cohesin and CTCF cooperate in regulating gene expression. Chromosome Res 17, 201–214, doi:10.1007/s10577-008-9017-7 (2009).

8 Hadjur, S. et al. Cohesins form chromosomal cis-interactions at the developmentally regulated IFNG locus. Nature 460, 410–U130, doi:10.1038/nature08079 (2009).

9 Kagey, M. H. et al. Mediator and cohesin connect gene expression and chromatin architecture. Nature 467, 430–435, doi:10.1038/nature09380 (2010).

10 Ciosk, R. et al. Cohesin’s binding to chromosomes depends on a separate complex consisting of Scc2 and Scc4 proteins. Molecular Cell 5, 243–254, doi:Doi 10.1016/S1097-2765(00)80420-7 (2000).

11 Gillespie, P. J. & Hirano, T. Scc2 couples replication licensing to sister chromatid cohesion in Xenopus egg extracts. Curr Biol 14, 1598–1603, doi:10.1016/j.cub.2004.07.053 (2004).

12 Watrin, E. et al. Human Scc4 is required for cohesin binding to chromatin, sister-chromatid cohesion, and mitotic progression. Curr Biol 16, 863–874, doi:10.1016/j.cub.2006.03.049 (2006).

13 Gandhi, R., Gillespie, P. J. & Hirano, T. Human Wapl is a cohesin-binding protein that promotes sister-chromatid resolution in mitotic prophase. Current Biology 16, 2406–2417, doi:10.1016/j.cub.2006.10.061 (2006).

14 Kueng, S. et al. Wapl controls the dynamic association of cohesin with chromatin. Cell 127, 955–967, doi:10.1016/j.cell.2006.09.040 (2006).

15 Tedeschi, A. et al. Wapl is an essential regulator of chromatin structure and chromosome segregation. Nature 501, 564-+, doi:10.1038/nature12471 (2013).

16 Arumugam, P. et al. ATP hydrolysis is required for cohesin’s association with chromosomes. Current Biology 13, 1941–1953, doi:10.1016/j.cub.2003.10.036 (2003).

17 Weitzer, S., Lehane, C. & Uhlmann, F. A model for ATP hydrolysis-dependent binding of cohesin to DNA. Curr Biol 13, 1930–1940 (2003).

18 Murayama, Y. & Uhlmann, F. Biochemical reconstitution of topological DNA binding by the cohesin ring. Nature 505, 367–371, doi:10.1038/nature12867 (2014).

19 Murayama, Y. & Uhlmann, F. DNA Entry into and Exit out of the Cohesin Ring by an Interlocking Gate Mechanism. Cell 163, 1628–1640, doi:10.1016/j.cell.2015.11.030 (2015).

20 Camdere, G., Guacci, V., Stricklin, J. & Koshland, D. The ATPases of cohesin interface with regulators to modulate cohesin-mediated DNA tethering. Elife 4, doi:10.7554/eLife.11315 (2015).

21 Yu, H. Magic Acts with the Cohesin Ring. Mol Cell 61, 489–491, doi:10.1016/j.molcel.2016.02.003 (2016).

22 Carretero, M., Ruiz-Torres, M., Rodriguez-Corsino, M., Barthelemy, I. & Losada, A. Pds5B is required for cohesion establishment and Aurora B accumulation at centromeres. Embo J 32, 2938–2949, doi:10.1038/emboj.2013.230 (2013).

23 Nishiyama, T. et al. Sororin Mediates Sister Chromatid Cohesion by Antagonizing Wapl. Cell 143, 737–749, doi:10.1016/j.cell.2010.10.031 (2010).

24 Hou, F. J. & Zou, H. Two human orthologues of Eco1/Ctf7 acetyltransferases are both required for proper sister-chromatid cohesion. Mol Biol Cell 16, 3908–3918, doi:10.1091/mbc.E04-12-1063 (2005).

25 Lafont, A. L., Song, J. & Rankin, S. Sororin cooperates with the acetyltransferase Eco2 to ensure DNA replication-dependent sister chromatid cohesion. Proc Natl Acad Sci U S A 107, 20364–20369, doi:10.1073/pnas.1011069107 (2010).

26 Whelan, G. et al. Cohesin acetyltransferase Esco2 is a cell viability factor and is required for cohesion in pericentric heterochromatin. Embo J 31, 71–82, doi:10.1038/emboj.2011.381 (2012).

27 Minamino, M. et al. Esco1 Acetylates Cohesin via a Mechanism Different from That of Esco2. Current Biology 25, 1694–1706, doi:10.1016/j.cub.2015.05.017 (2015).

28 Higashi, T. L. et al. The prereplication complex recruits XEco2 to chromatin to promote cohesin acetylation in Xenopus egg extracts. Curr Biol 22, 977–988, doi:10.1016/j.cub.2012.04.013 (2012).

29 Song, J. et al. Cohesin acetylation promotes sister chromatid cohesion only in association with the replication machinery. J Biol Chem 287, 34325–34336, doi:10.1074/jbc.M112.400192 (2012).

30 Minamino, M. et al. Temporal Regulation of ESCO2 Degradation by the MCM Complex, the CUL4-DDB1-VPRBP Complex, and the Anaphase-Promoting Complex. Curr Biol 28, 2665–2672 e2665, doi:10.1016/j.cub.2018.06.037 (2018).

31 Ivanov, M. P. et al. The replicative helicase MCM recruits cohesin acetyltransferase ESCO2 to mediate centromeric sister chromatid cohesion. Embo J 37, doi:10.15252/embj.201797150 (2018).

32 Price, J. C. et al. Sequencing of candidate chromosome instability genes in endometrial cancers reveals somatic mutations in ESCO1, CHTF18, and MRE11A. PLoS One 8, e63313, doi:10.1371/journal.pone.0063313 (2014).

33 Schule, B., Oviedo, A., Johnston, K., Pai, S. & Francke, U. Inactivating mutations in ESCO2 cause SC phocomelia and Roberts syndrome: No phenotype-genotype correlation. Am J Hum Genet 77, 1117–1128, doi:Doi 10.1086/498695 (2005).

34 Vega, H. et al. Roberts syndrome is caused by mutations in ESCO2, a human homolog of yeast ECO1 that is essential for the establishment of sister chromatid cohesion. Nat Genet 37, 468–470, doi:10.1038/ng1548 (2005).

35 Gordillo, M. et al. The molecular mechanism underlying Roberts syndrome involves loss of ESCO2 acetyltransferase activity. Hum Mol Genet 17, 2172–2180, doi:10.1093/hmg/ddn116 (2008).

36 Van Den Berg, D. J. & Francke, U. Roberts syndrome: a review of 100 cases and a new rating system for severity. Am J Med Genet 47, 1104–1123, doi:10.1002/ajmg.1320470735 (1993).

37 Kouznetsova, E. et al. Sister Chromatid Cohesion Establishment Factor ESCO1 Operates by Substrate-Assisted Catalysis. Structure 24, 789–796, doi:10.1016/j.str.2016.03.021 (2016).

38 Rivera-Colon, Y., Maguire, A., Liszczak, G. P., Olia, A. S. & Marmorstein, R. Molecular Basis for Cohesin Acetylation by Establishment of Sister Chromatid Cohesion N-Acetyltransferase ESCO1. J Biol Chem 291, 26468–26477, doi:10.1074/jbc.M116.752220 (2016).

39 Chao, W. C. et al. Structural Basis of Eco1-Mediated Cohesin Acetylation. Sci Rep 7, 44313, doi:10.1038/srep44313 (2017).

40 Salah Ud-Din, A. I., Tikhomirova, A. & Roujeinikova, A. Structure and Functional Diversity of GCN5-Related N-Acetyltransferases (GNAT). Int J Mol Sci 17, doi:10.3390/ijms17071018 (2016).

41 Ladurner, R. et al. Cohesin’s ATPase activity couples cohesin loading onto DNA with Smc3 acetylation. Curr Biol 24, 2228–2237, doi:10.1016/j.cub.2014.08.011 (2014).

42 Gligoris, T. G. et al. Closing the cohesin ring: structure and function of its Smc3-kleisin interface. Science 346, 963–967, doi:10.1126/science.1256917 (2014).

43 Cheng, K. C., Liao, J. N. & Lyu, P. C. Crystal structure of the dopamine N-acetyltransferase-acetyl-CoA complex provides insights into the catalytic mechanism. Biochem J 446, 395–404, doi:10.1042/BJ20120520 (2012).

44 Fitzgerald, D. J. et al. Protein complex expression by using multigene baculoviral vectors. Nat Methods 3, 1021–1032, doi:10.1038/nmeth983 (2006).

45 Trowitzsch, S., Bieniossek, C., Nie, Y., Garzoni, F. & Berger, I. New baculovirus expression tools for recombinant protein complex production. J Struct Biol 172, 45–54, doi:10.1016/j.jsb.2010.02.010 (2010).

46 Kabsch, W. Xds. Acta Crystallogr D Biol Crystallogr 66, 125–132, doi:10.1107/S0907444909047337 (2010).

47 Emsley, P., Lohkamp, B., Scott, W. G. & Cowtan, K. Features and development of Coot. Acta Crystallogr D Biol Crystallogr 66, 486–501, doi:10.1107/S0907444910007493 (2010).

48 Adams, P. D. et al. PHENIX: a comprehensive Python-based system for macromolecular structure solution. Acta Crystallogr D Biol Crystallogr 66, 213–221, doi:10.1107/S0907444909052925 (2010).

49 Rodriguez, C. I. et al. High-efficiency deleter mice show that FLPe is an alternative to Cre-loxP. Nature Genetics 25, 139–140, doi:Doi 10.1038/75973 (2000).

50 Ruzankina, Y. et al. Deletion of the developmentally essential gene ATR in adult mice leads to age-related phenotypes and stem cell loss. Cell Stem Cell 1, 113–126, doi:10.1016/j.stem.2007.03.002 (2007).

51 Mendez, J. & Stillman, B. Chromatin association of human origin recognition complex, cdc6, and minichromosome maintenance proteins during the cell cycle: assembly of prereplication complexes in late mitosis. Mol Cell Biol 20, 8602–8612 (2000).

52 Gouet, P., Courcelle, E., Stuart, D. I. & Metoz, F. ESPript: analysis of multiple sequence alignments in PostScript. Bioinformatics 15, 305–308 (1999).

